# Win-concurrent sensory cues can promote riskier choice

**DOI:** 10.1101/323345

**Authors:** Mariya V. Cherkasova, Luke Clark, Jason J.S. Barton, Michael Schulzer, Mahsa Shafiee, Alan Kingstone, A. Jon Stoessl, Catharine A. Winstanley

## Abstract

Reward-related stimuli can potently influence behaviour; for example, exposure to drug-paired cues can trigger drug use and relapse in people with addictions. Psychological mechanisms that generate such outcomes likely include cue-induced cravings and attentional biases. Recent animal data suggest another candidate mechanism: reward-paired cues can enhance risky decision making, yet whether this translates to humans is unknown. Here, we examined whether sensory reward-paired cues alter decision making under uncertainty and risk, as measured respectively by the Iowa Gambling Task and a two-choice lottery task. In the cued version of both tasks, gain feedback was augmented with reward-concurrent audiovisual stimuli. Healthy human volunteers (53 males, 78 females) performed each task once, one with and the other without cues (cued IGT/uncued VGT: n = 63; uncued IGT/cued VGT: n = 68), with concurrent eye-tracking. Reward-paired cues did not affect choice on the Iowa Gambling Task. On the two-choice lottery task, the cued group displayed riskier choice and reduced sensitivity to probability information. The cued condition was associated with reduced eye fixations on probability information shown on the screen and greater pupil dilation related to decision and reward anticipation. This pupil effect was unrelated to the risk-promoting effects of cues: the degree of pupil dilation for risky versus risk-averse choices did not differ as a function of cues. Taken together, our data show that sensory reward cues can promote riskier decisions and have additional and distinct effects on arousal.

**SIGNIFICANCE STATEMENT:** Animal data suggest that reward-paired cues can promote maladaptive reward-seeking by biasing cost-benefit decision making. Whether this finding translates to humans is unknown. We examined the effects of salient reward-paired audio-visual cues on decision making under risk and uncertainty in human volunteers. Cues had risk-promoting effects on a risky choice task and independently increased task-related arousal as measured by pupil dilation. By demonstrating risk-promoting effects of cues in human participants, our data identify a mechanism whereby cue reactivity could translate into maladaptive behavioural outcomes in people with addictions.

## Introduction

Reward-linked environmental stimuli, commonly described in psychological studies as “cues”, can potently influence behaviour. In addicted individuals, exposure to cues such as drug paraphernalia can trigger cravings, drug use and relapse (Childress et al., 1993). Cues may likewise play a role in supporting behavioural addictions such as Gambling Disorder. Electronic gambling machines, which feature complex and salient audio-visual cues, are associated with some of the highest rates of disordered gambling (Dowling et al., 2005).

The incentive sensitization theory of addiction posits that, through Pavlovian associations with primary rewards (e.g. intoxication or thrill of winning), cues acquire incentive salience and come to act as motivational magnets (Robinson and Berridge, 1993). By eliciting cravings and capturing attention (Carter and Tiffany, 1999; Field and Cox, 2008), cues might help define behavioural goals, thus encouraging pursuit of the addiction. However, cue-elicited cravings and attentional biases do not explain how these goals translate into the series of actions required to achieve them, particularly in the face of other conflicting goals such as abstinence. When the choice is made to engage in addictive behaviour, the benefits may be judged to outweigh the costs. Indeed, impairments in cost-benefit decision making are well documented in substance and behavioural addictions (Grant et al., 2000; Bechara et al., 2001; Hanson et al., 2008; Kovács et al., 2017). Here, we consider whether cues influence cost-benefit decision making, thereby providing a candidate mechanism that enables transition from cue-elicited motivational states to the maladaptive actions that support addiction.

Recent rodent data support this hypothesis. Pairing food rewards with audio-visual cues increased risky choice on a rodent gambling task modelled after the Iowa Gambling Task (IGT) in a dopamine D3 receptor-dependent manner (Barrus and Winstanley, 2016). To our knowledge, no such data are available in humans. In simulated gambling paradigms, gambling-related sensory cues have been found to increase play enjoyment and arousal, as well as to distort estimates of earned profits (Dixon et al., 2010; Dixon et al., 2014; Dixon et al., 2015); however, no effects on choice *per se* have been reported. One study directly examining the effects of lighting and casino sound on IGT performance found that choices were unaffected by these cues, though their presence did elevate mood and abolish the slowing of response times on trials following losses (Brevers et al., 2015). Other evidence suggests that presentation of aversively conditioned stimuli and past reward primes can modulate risk preferences (Guitart-Masip et al., 2010; Ludvig et al., 2015).

We therefore examined the effects of casino-inspired sensory reward cues on decision making in human participants using two laboratory tasks. We chose the IGT (Bechara et al., 1994), as most analogous to the rodent task and given the considerable evidence of impairments on this task in substance use and gambling disorders (Grant et al., 2000; Bechara et al., 2001; Hanson et al., 2008; Kovács et al., 2017). We also used a two-choice lottery task, to which we refer as the Vancouver Gambling Task (Sharp et al., 2012; Sharp et al., 2013), to enable a behavioural economic analysis of risk preferences. Versions of both tasks were created in which reward feedback was either accompanied or unaccompanied by audio-visual cues. We hypothesized that these cues would have risk-promoting effects in both tasks.

In addition to decision making, we explored: 1) the pattern of eye fixations during choice to help elucidate the mechanisms of cue-induced behavioural effects; 2) pupil dilation as a proxy of arousal. Changes in pupil size are closely coupled to noradrenaline signaling (Murphy et al., 2014; Joshi et al., 2016) and co-vary with a number of psychological variables, including decision making (Einhäuser et al., 2010; Preuschoff et al., 2011; de Gee et al., 2017), though it is unclear to which decision variables the pupil responds (Einhauser, 2017). In light of the previous finding that slot machine sounds increased arousal (Dixon et al., 2014), we hypothesized greater pupil dilation in the cued condition.

## Materials and Methods

### Participants

131 healthy human volunteers recruited from the community took part in the study (males: n = 53, females: n = 78, mean age = 25.65 ± 8.28). The sample size was based on the assumption of a medium effect size for the effect of sensory cues on decision making. According to GPower, 128 participants are required to have the power of .8 to detect a medium effect at p = .05 in an ANOVA. Participants were required to have normal or corrected-to-normal vision and hearing. Although these were the only inclusion criteria, detailed self-report data were collected from the participants regarding their medication and substance use using Module E from the Structured Clinical Interview for DSM-IV disorders (SCID-IV) (First et al., 2002). Five participants reported ongoing use of psychotropic medications: escitalopram for major depression (n = 1); stimulants for attention deficit hyperactivity disorder (n = 4). Two participants met criteria for substance dependence. Because excluding the data from these participants did not change the significance of the findings, we report the results from the entire sample. The study was conducted in accordance with institutional guidelines and the Declaration of Helsinki, and was approved by the Research Ethics Board of the University of British Columbia. Participants gave written informed consent. Compensation for the study corresponded to the bonus amount earned on the tasks.

### Procedure

Participants were randomly assigned to two groups. Group 1 (n = 63) performed the IGT with the sensory cues (henceforth ‘cued’) and the VGT without the sensory cues (henceforth ‘uncued’). Group 2 (n = 68) performed the uncued IGT and the cued VGT. The order of tasks (IGT first vs. VGT first) was randomized and counterbalanced between groups. The two groups did not differ significantly in terms of age or gender composition (p_s_ ≥ 0.62).

Eye fixations and pupil size data were obtained using the EyeLink 1000 infrared pupil tracker with a sampling rate of 1000 Hz and resolution of 0.01° of visual angle (SR Research Ltd., Mississauga, Ontario). Most participants (n = 85) were tested on the EyeLink 1000 Tower system. The remaining participants were tested using the EyeLink 1000 Desktop system, either because they required glasses or because the Tower system was unavailable. Participants were tested in one of two labs, in which the apparatus differed slightly. The majority (n = 115) were tested in a slightly dimmed lab (illumination = 80 lux) with the eyes positioned at the distance of 60cm from a 22’’ monitor. Sixteen participants were tested using the Desktop system in a lab without a dimmer, so the testing room was dark (1 lux); the monitor was 17’’ in size, so the viewing distance was adjusted accordingly (47 cm) to preserve stimulus size in visual angle. Prior to each task, a 5-point calibration was performed. Stimulus presentation and data collection were controlled via scripts developed using the eye tracker’s proprietary software Experiment Builder.

### Iowa Gambling Task

The IGT presented participants a choice between 4 decks of cards. They were informed that with each selected card they could win or lose money and that some decks were more advantageous than others. Unlike the standard IGT, in which both gains and losses can occur on any trial, the current version presented either a net gain or a net loss. This simplified outcome structure facilitated congruency between reward cues and net outcome value, while also achieving closer correspondence to the rodent gambling task (Barrus and Winstanley, 2016), in which gains and losses are not simultaneous. Two of the decks were high-risk and high-reward decks, resulting in larger gains on successful trials ($100), but also in larger and/or more frequent losses: 10% chance of losing $1150 for one of the decks and 50% change of losing either $50, $100, $150, $200 or $250 for the other deck. The remaining two decks were low-risk and low-reward, resulting in smaller gains ($50) on successful trials and either no loss or a smaller or infrequent losses on unsuccessful trials: 50% chance of a $0 outcome for one of the decks and 10% change of losing $150 for the other deck. Over time, choosing from the low-risk low-reward decks typically yields a cumulative gain, making this the optimal strategy. Participants were told that they would receive 10% of the amount accumulated over the 100 trials. They started with a bank of $2000, corresponding to $20 (Canadian) in actual money. Each trial comprised the a decision phase, during which the participant chose from one of the four decks (no time limit imposed), followed by a feedback phase (3500ms), during which subjects were shown how much they had won or lost on that trial, with their total earnings updated at the top of the display. The task could yield negative earnings, but these were counted as 0 for the purposes of the bonus payout. Participants wore the eye tracker during the IGT but its temporal structure was not suitable for eye tracking, due to the absence of an inter-trial interval to establish a trial baseline.

In the uncued IGT, feedback regarding gains and losses was given numerically. In the cued version, feedback about gains (but not losses) was augmented by audiovisual reward cues. Gains ($50 for safer decks and $100 for riskier decks) were represented by images of stacks of Canadian $10 bills accompanied by casino jingles (Figure 1A). The $100 and $50 images were identical in luminance and color (luminance: 38.75 cd/m^2^; color: .226 u’; .482 v’) and differed only in the number of bills in the stack. The auditory jingles were taken from a casino sound library and edited to conform to the temporal structure of the task. The jingle was longer (1200 ms for $50 versus 2700ms for $100), louder (44 dB for $50 versus 52dB for $100) and more complex (greater variation in pitch and tempo) for the larger win.

**Figure 1:**
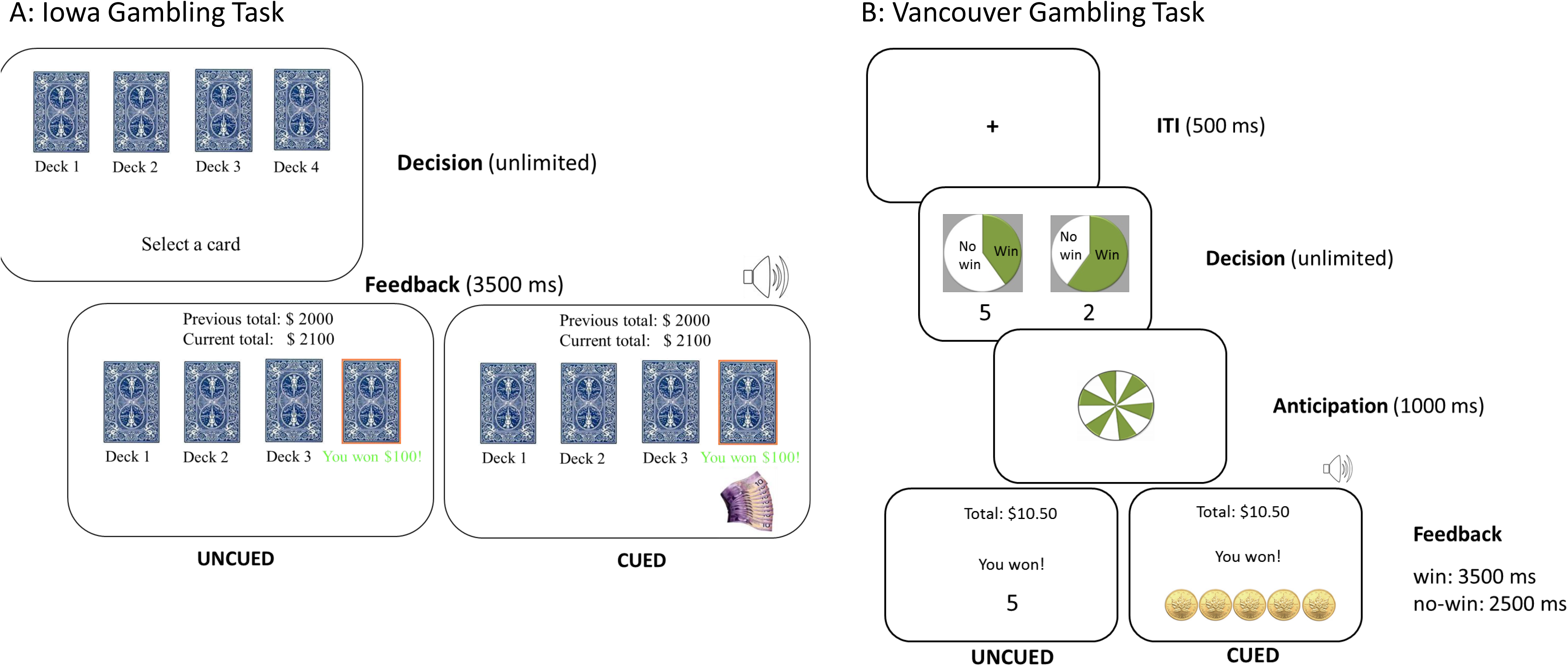
Iowa and Vancouver Gambling Tasks.

### Two Choice Lottery Task

The two choice lottery task, which we label the Vancouver Gambling Task (VGT), consistent with the previous published studies, assesses willingness to take risks at different combinations of reward probability and magnitude. As such, it permits to model the impact of the reward’s expected value (EV), its probability and its magnitude on risk attitudes.

Participants made a choice between two prospects on every trial. One prospect featured a larger but less probable gain, while the other featured a smaller and more probable gain. There were 10 unique prospect pairs, each repeated 10 times for the total of 100 trials. These 10 pairs formed a continuum of relative EVs of the options, ranging from pairs that highly favored the ‘safer’ choice (i.e. the smaller but more probable prospect) to pairs that highly favored the ‘riskier’ choice (i.e. the larger but less probable prospect). Thus, each pair was associated with a unique Expected Value Ratio (EV-ratio) calculated as [EV(safe) – EV(risky)]/mean(EV(safe), EV(risky)) as per (Sharp et al., 2012).

Each trial started with a 500ms inter-trial interval, displaying a fixation cross in the center of the screen. Next came the decision phase, in which participants were shown the two prospects, and could take as much time as required to choose one. Probabilities for each prospect were represented as pie charts, with a green sector representing the odds of winning, which always summed to 100%: 20% vs. 80%, 30% vs. 70%, 40% vs. 60%. The location (left vs. right) of the higher probability (safer) vs. lower probability (riskier) option was randomized across trials for every testing session, with the constraint that safer and riskier options appeared an equal number of times on the left and on the right. Gain magnitudes were represented using numerals beneath the pie charts, indicating the number of tokens that could be won (1, 2, 3, 4 or 5 tokens), with each token worth 10 Canadian cents in actual money. The decision phase was followed by a 1000 ms anticipation phase with a spinning roulette display.

On each trial, the participant either won the reward depicted in the chosen gamble or received nothing: no losses occurred. In the uncued VGT gain feedback was delivered using numerals, without sound accompaniment. In the cued VGT, the gains were represented by images of coins accompanied by casino jingles (Figure 1B). These visual and auditory cues scaled in sensory intensity and complexity with gain magnitude. The visual enhancement was as follows: 1 token was represented as a static two-dimensional image of 1 gold coin; 2 tokens as 2 static two-dimensional gold coins with a sparkle (static luminance enhancement); 3 tokens as 3 static three-dimensional gold coins with 3 sparkles (static luminance and depth enhancement); 4 tokens as 4 three-dimensional gold coins with a sparkle running along the circumference of each coin (dynamic luminance and depth enhancement); 5 tokens as 5 three-dimensional spinning gold coins with a sparkle running along the circumference of each coin (dynamic luminance, depth and motion enhancement). Despite the visual enhancement, the visual stimuli for the different reward magnitudes were not substantially different in terms of the overall average luminance (1 coin: 100.79 cd/m^2^; 2 coins: 101.54 cd/m^2^; 3 coins: 98.93 cd/m^2^; 4 coins: 94.37 cd/m^2^; 5 coins: 101.18 cd/m^2^) and color of the image (1 coin: .215 v’, .500 u’; 2 coins: .240 v’, .531 u’; 3 coins: .228 v’, .525 u’; 4 coins: .186 v’, .387u’; 5 coins: .229 v’, .528 u’); the 3-dimensional images were slightly lower in luminance because of the shading. The average luminance of the uncued feedback image was 102.44 cd/m^2^. The auditory enhancement again consisted of sounds taken from a casino library and edited to conform to the temporal structure of the task. The tunes accompanying the rewards progressively increased in duration (1200 to 2700 ms), loudness (44 to 52dB) and complexity (variation in tempo and pitch) as the reward magnitude increased from 1 token to 5 tokens. The cued VGT auditory stimulus accompanying a 2 token win was the same as the auditory stimulus accompanying the smaller ($50) gain on the cued IGT; the stimulus accompanying a 5 token win on the VGT was the same as the one accompanying larger ($100) gains on the cued IGT. To ensure that participants understood the task, they were given 5 practice trials before the start of the 100 test trials. At the end of every 20 trials they were offered a break.

Following IGT and VGT, participants additionally performed a Pavlovian-instrumental transfer paradigm modelled after (Garofalo and di Pellegrino, 2015): data not shown.

### Analyses

#### Decision making

Statistical analyses were performed using the lme4 package in R (Bates et al., 2015); R syntax for the models is provided. The effect of cues on the number of advantageous choices over the progression of the 5 blocks was analyzed using a linear mixed effects model with a logistic link (glmer function) predicting optimal vs. risky choice on a trial-by-trial basis as a function of sensory cues in interaction with task block (to evaluate changes in learning rate as a function of feedback type). The model included task order (VGT before IGT or vice versa) as an additional fixed factor and random intercepts for participants.

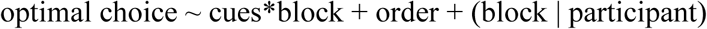

In light of the previously reported abolition of post-error slowing in the presence of casino lighting and sound (Brevers et al., 2015), we tested the effect of sensory cues on reaction times following losses versus wins. This was achieved by modelling response time as a function of prior outcome (fixed effect) in interaction with sensory cues (fixed effect); task order was included as an additional fixed factor and random intercepts were modelled for participants. Response times were log-transformed for analysis. Given that response times decreased as a function of block (b= 0.05, SE=0.0013, t=37.72, p < 0.0005), and the slopes of this effect varied across participants, the model also included random slopes for blocks.

The effect of sensory cues on VGT performance was analyzed using linear mixed effects models with a logistic link (glmer function). The first modelled choice of a higher-probability (safer) prospect versus a lower-probability (riskier) prospect on a trial-by-trial basis as a function of sensory cues (fixed effect) in interaction with EV-ratio (fixed effect), with task order included as an additional fixed factor and random intercepts for participants. Because participants made riskier choices with successive trial repetitions (b = 0.92, SE = 0.10, t = 9.69, p < 0.0005), and the slope of this effect varied across participants, the model also included random slopes for trial repetition.

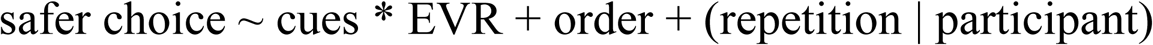

The second model considered the influence of sensory cues on evaluation of reward probabilities and magnitudes in determining safe vs. risky choice. Because probabilities and magnitudes of the two alternatives were evaluated relative to each other, we performed isometric log ratio transformations on probability and magnitude pairs to derive a single value for each representing, respectively, relative probabilities and relative magnitudes of the alternatives in each trial. Relative probabilities and relative magnitudes of alternatives were modelled as fixed effect terms in interaction with sensory cues.

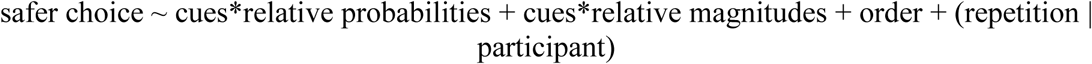

Finally, because the sensory cue manipulation only affected wins, we modelled choice as a function of prior trial’s outcome (win vs. 0-outcome) in interaction with sensory cues. The random effect structure was the same as in the first model, and task order was modelled as a fixed factor.

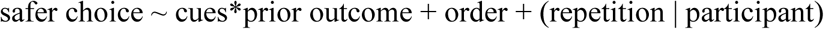

#### Gaze fixations

We examined the effect of sensory cues on attention allocated to probability and magnitude information on the screen as indexed by fixations. For each trial, we quantified percent of total time of the decision phase spent looking at each of 3 most fixated interest areas (IA): 1) the two probability information zones, 2) the two magnitude information zones, and 3) the screen centre (Figure 2D). Rectangular IAs were defined based on aggregate fixation duration heat maps using the EyeLink Data Viewer software; the same IAs were used for both cued and uncued data. We analyzed the extracted % fixation duration values using a linear mixed effects model (lmer function) with sensory cues and IA as fixed effects in interaction; random intercepts and slopes with respect to IA were modeled for participants.

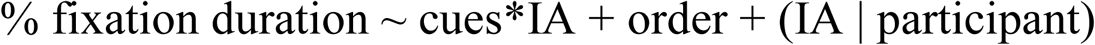

**Figure 2:**
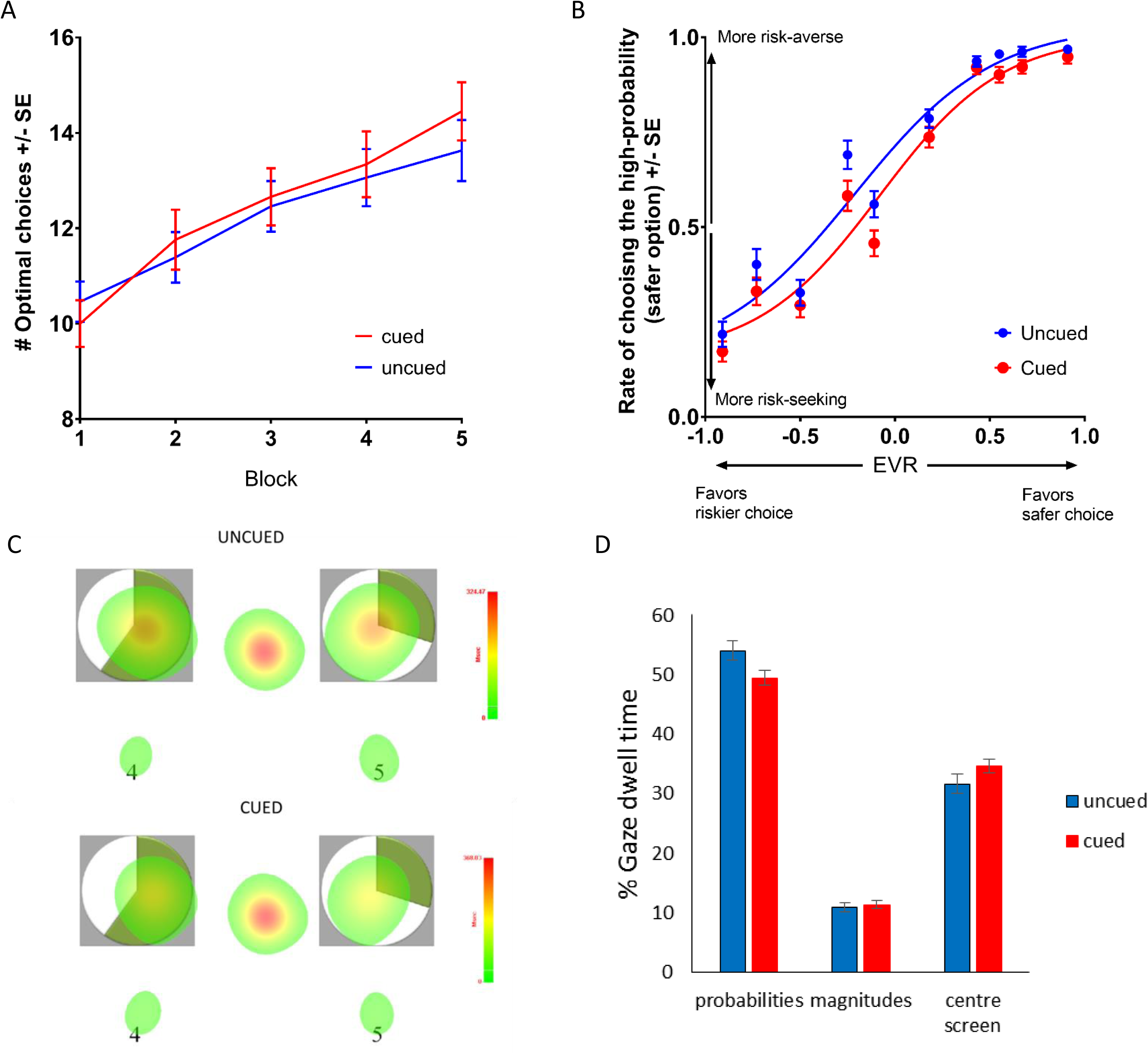
The effect of sensory reward cues on Iowa and Vancouver Gambling Task performance. A) Number of advantageous choices on the IGT as a function of block. B) Rate of risk-averse choices as a function of Expected Value Ratio. EVR = (EVsafe_choice – EVrisky choice)/mean(EVsafe_choice, EVrisky_choice). The curves are fitted using a 4-parameter logistic function. The downward shift of the curve for VGT with sensory cues indicates higher rate of choosing the riskier prospect with a higher potential payout independent of EV. This risk-promoting effect is driven by diminished influence of probability information on choice. C) Fixation heat maps representing fixation durations during the decision phase. D) Group averages of fixation durations during the decision phase (*p <.05).

#### Pupillometry

We focused on the subset of 85 participants (43 completing uncued VGT; 42 completing cued VGT) whose data were collected using the EyeLink 1000 Tower system. Because the EyeLink 1000 in centroid mode measures pupil size in angular units, which represent the area subtended by the pupil from the point of view of the camera (Hayes and Petrov, 2016b), pupil size measures can differ depending on the eye-to-camera distance. This distance differs for Tower and Desktop system and can vary for the Desktop system depending on the experimental layout.

Pupil time-series for the VGT were extracted for each participant and processed using Matlab scripts. First, a linear interpolation was performed over all samples occurring during blinks, using 100ms prior to and after each blink as start and endpoints of the interpolations. Next, the pupil time series were smoothed using a 2^nd^ order Butterworth low pass filter with the cut-off frequency of 4 Hz and down-sampled to 100Hz. The pupil time-series for each trial was then transformed into a time-series representing modulation in pupil area: percent change in pupil area (p) was computed at each time point in the series (t) with respect to baseline [(p(t)-baseline)/baseline*100]. Baseline was taken to be the average pupil area during the first 200ms of the decision phase: as pupillary response lags behind the stimulus by ~ 400 ms, peaking about 1-2 seconds post-stimulus (Partala and Surakka, 2003; Clayton et al., 2004), pupil area during the initial 200ms of the decision phase likely reflects effects from the inter-trial interval and not from the stimuli shown in the decision phase. Given that the duration of the decision phase was determined by the participant’s response time, the pupil modulation series was time-locked to the end (last 500ms) of the decision phase, not its start. Also, because the phasic responses of noradrenergic neurons within the locus coeruleus (LC) appear to be time-locked to the behavioural response rather than the stimulus (Clayton et al., 2004), a period linked to the end of the decision phase is likely most relevant.

We then used the pupil modulation time-series to calculate the area under the curve (AUC) of pupil response for each trial in every participant to be used as the outcome measure in the statistical analyses (Figure 3). The AUC outcome measure combined decision and anticipation phases of the task and excluded the pupil responses to the feedback phase because of differences in the visual stimulus for the cued and uncued VGT. For the decision-related pupil response we included data from an interval from 500ms before the end of the decision phase to the 400ms after its end, given the 400ms lag in pupil response. For the anticipation phase the interval spanned from 400ms to 1400ms after the end of the decision phase. We performed statistical analysis of AUC pupil dilation using linear mixed effects models (lmer function); log-transformed AUC values were used to ensure scale similarity with other variables in the model. As we wanted to test a) whether sensory feedback was associated with increased pupil dilation and b) whether this increase was associated with any cue-induced shift in risky choice, we modelled AUC pupil dilation as a function of sensory cues (fixed effect) in interaction with choice (risky vs. safe, fixed factor). The model also included the outcome of the prior trial in interaction with the sensory cues: because cues only accompanied wins and not 0-outcomes, effects of cues should be preferentially linked to these outcomes.

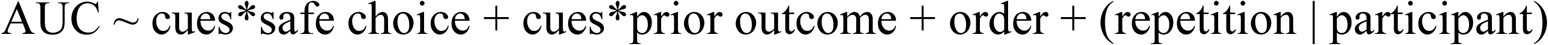

**Figure 3.**
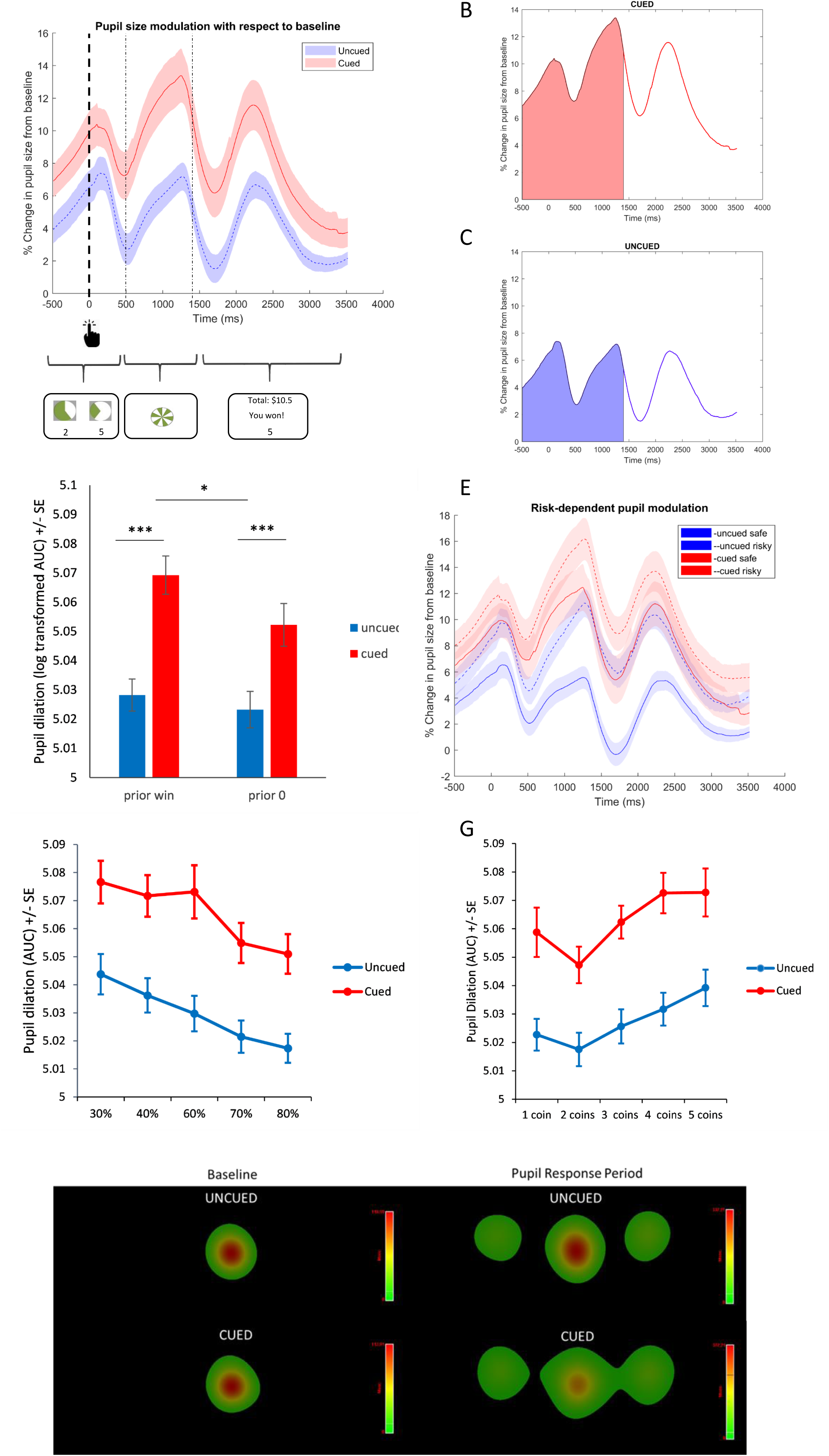
Pupil dilation during Vancouver Gambling Task. A) Pupil modulation time courses for VGT with and without sensory cues over the different trial epochs. Modulation is computed as % change from baseline in pupil size (measured as area subtended by the pupil in angular units from the point of view of the camera) over the course of last 500ms of the decision phase, the anticipation phase and the feedback phase. B) Area under the curve (AUC) measure of pupil dilation from baseline over the decision and anticipation periods for VGT with the sensory cues. C) AUC pupil dilation over the decision and anticipation periods without sensory cues. D) Pupil dilation with respect to baseline following wins and 0-outcomes for VGT with and without sensory cues. Pupil dilation is plotted as log-transformed AUC measure depicted in B. E) Pupil dilation with respect to baseline for safe (higher probability prospect, solid lines) vs. risky (lower probability prospect, dotted lines) choices. F) Pupil dilation with respect to baseline as a function of the chosen prospect’s probability. G) Pupil dilation with respect to baseline as a function of the chosen prospect’s magnitude. H) Heat maps of fixation durations during baseline (first 200ms of the decision phase) and the analyzed pupil response period (final 500ms of decision plus the anticipation phase). *p < 0.05; ***p < 0.0005.

Additional models were used to explore relationships between pupil dilation and prospect characteristics (probabilities and magnitudes), which were modelled as fixed effects in interaction with sensory cues. As the model considering probability and magnitude only of the chosen prospect better fit the data than the one considering probability and magnitude information for both prospects in the gamble (Akaike Information Criterion (AIC): ‐18511 and ‐18479, respectively), we report the results only for the model uniquely considering the chosen prospect characteristics. This is in keeping with the view that phasic LC activation reflects decision outcome (Aston-Jones and Cohen, 2005), and pupil dilation response should therefore primarily co-vary with the characteristics of the chosen prospect. Task order was considered as an additional fixed factor. Random intercepts and slopes with respect to trial repetition were modelled for participants. The latter was because of a progressive decrease in pupil response (b = 0.001, SE = 0.0003, t= 3.20, p <0.001) over the course of the task, the slope of which varied across participants.

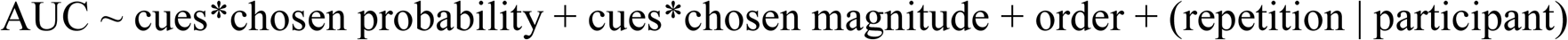

Finally, because pupil foreshortening error at eccentric eye positions can affect pupil size measurements (Hayes and Petrov, 2016b), and there is no validated pupil foreshortening correction algorithm that could be implemented with our data collected using the EyeLink Tower system, which was used to collect the majority of our data, we analyzed the pattern of eye fixations during the time interval pertaining to the pupillometry analysis. This was done in the same way as the fixation analysis for the decision phase, but using different IAs: left, right and centre. These IAs were defined based on the fixation heat map from the time period in question. Because fixation eccentricity (rather than the content of the fixated region) was the variable of interest in this analysis, we examined separately fixations to the left and to the right off centre.

## Results

### Decision making on the Iowa Gambling Task

Sensory cues did not have a significant effect on the number of advantageous choices on the IGT either on its own or in interaction with block (p_s_ < 0.26). Participants made more advantageous choices as they progressed through the task blocks (b = 0.35, SE = 0.05, z = 7.64, p < 0.0005), but the slope of this change did not differ as a function of cues (Figure 2A). As expected, the prior trial’s outcome predicted the choice of advantageous decks (b = .05, SE = .01, z = 3.53, p = 0.0004). There was also a significant interaction of cues with prior outcome (b = .05, SE = .02, z = 2.20, p = 0.03): participants performing the cued IGT were less likely to choose advantageous decks following large losses. In addition, participants who performed the IGT first showed a trend towards choosing less advantageously than the ones who performed it after the VGT (b = .23, SE = 0.13, z = 1.79, p = 0.07), but this task order effect did not interact with cues.

For response times (RTs), there was no significant effect of prior outcome (win vs. loss) or sensory cues, or interaction of these two terms (p_s_ ≥ 0.45).

### Decision making on the Vancouver Gambling Task

There was a significant main effect of sensory cues on choice (b = 0.58, SE = 0.22, z= 2.64, p = 0.008) without an interaction with EV-ratio (p = 0.55), indicating that choices were more risk-seeking in the presence of the sensory cues. This effect was independent of expected value (Figure 2B). The model considering the impact of sensory cues on the evaluation of probability and magnitude information revealed a significant interaction of sensory cues with the probability term (b = 0.68, SE = 0.28, z= 2.36, p = 0.02), but not with the magnitude term (p = .32): choices were less probability-driven in cued version than in the uncued version (cued: b = 3.88, SE = 0.33; uncued: b =4.04, SE = 0.35). Finally, although prior 0-outcomes precipitated riskier choices (b = 0.2, SE = 0.04, z = 5.26; p < 0.0005), this did not interact significantly with sensory cues (p = 0.99).

### Fixations

The analysis of gaze fixations during decision making provided additional evidence of decreased consideration of probability information in the cued VGT (Figure 2C). There was a significant interaction of cues with IA (b = 0.03, SE = 0.02, t = 1.97; p = 0.05). During the decision phase, the cued group spent a smaller proportion of time fixating on the probability pie charts (b = 0.02, SE = 0.01, t = 2.26; p = 0.03), but did not differ significantly from the uncued group on the time spent fixating the other IAs (p_s_ ≥ 0.1).

### Pupillometry

We first confirmed the behavioural effects of sensory cues on VGT in the pupillometry subsample. The main effect of the sensory cues remained significant (b = 0.58, SE = 0.22, z= 2.64, p = 0.008), as was the interaction of cues with the probability term (b = 0.69, SE = 0.34, z= 2.05, p = 0.04).

A model predicting AUC pupil dilation as a function of sensory cues in interaction with choice (safe vs. risky) and with prior outcome (wins vs. 0-outcomes) revealed a significant effect of cues (b = 0.027, SE = 0.0085, t= 3.18, p =0.002), with greater pupil dilation in the cued VGT (Figure 3A, B, C). Relative to prior 0-outcomes, prior wins predicted greater pupil dilation in the subsequent trial (b = 0.014, SE = 0.002, t= 6.44, p < 0.0005), and this effect interacted with sensory cues (b = 0.012, SE = 0.004, t = 3.5, p = 0.0005): the potentiation of pupil dilation on trials following wins (relative to those following 0-outcomes) was amplified by sensory cues (Figure 3D). Finally, there was a significant effect of choice (b = 0.016, SE = 0.002, t= 8.11, p <0.0005), with risky choices associated with greater pupil dilation in both cued and uncued versions (Figure 3E). This effect did not interact with cues (p=.68). There was no significant difference in baseline pupil size between cued and uncued VGT (p = 0.31).

We next explored whether probability and magnitude of the chosen prospect modulated the amplification of pupil dilation by sensory cues. There were significant and opposite effects of probability (b = 0.057, SE = 0.019, t= 4.14, p <0.0005) and magnitude (b = 0.002, SE = 0.0009, t= 2.22, p <0.03) on pupil dilation: choosing either less likely and more rewarding prospects was associated with greater pupil dilation (Figure 3F, G). These effects were present over and above the effects of risky choice *per se*, as they were apparent in the presence of the safe vs. risky choice variable in the model (b = 0.01, SE = 0.005, t= 1.97, p = 0.05). Neither the probability nor the magnitude term interacted with sensory cues to predict pupil dilation, although there was a trend for the chosen reward size to modulate pupil response more in the absence of the cues (b = 0.003, SE = 0.002, t= 1.74, p = 0.08).

The analysis of gaze fixations in the cued vs. uncued VGT focusing on the time frame of the pupillometry analysis did not reveal any significant differences in fixation patterns (p_s_ ≥ 0.21). It is unlikely that the small non-significant differences in gaze eccentricity during the pupil response period that we considered drove the differences in pupil dilation between cued and uncued conditions: if anything, gaze tended to be more eccentric in the cued VGT (Figure 3H), which would be associated with larger pupil foreshortening error and consequently smaller measured pupil area, as the pupil assumes a more elliptical shape with more eccentric gaze.

## Discussion

Our data directly demonstrate for the first time that reward-concurrent sensory cues can promote risky choice in human subjects. While cue reactivity literature focuses on cues that represent incentives or predict rewards, here the cues accompanied rewards, rather than rather than being positioned to predict them, in keeping with the way such cues are used in commercial gambling products.

The VGT enabled decomposition of risky choice in terms of sensitivity to expected value, probability and magnitude of the prospects. We found that the risk enhancement produced by sensory cues was independent of expected value, but reflected a decreased influence of reward probability on choice. This finding was further supported by the pattern of eye movements in the course of decision making, with proportionally less time spent fixating on the probability information depicted on the screen. Heavy reliance on probability information leads to risk-averse performance on the VGT (high rates of choosing the higher probability prospects); sensory cues decreased this tendency. The mechanism(s) through which cues shift the emphasis away from probability information remains unclear. One possibility is that augmenting gains with sensory feedback enhances the memorability of gains, which then biases the perceived probability of winning via the availability heuristic. We did not have a subjective measure of perceived gain probabilities, but a conceptually similar effect has been previously reported in gambling research. In electronic gambling machines, “losses disguised as wins” – net loss events that are accompanied by the ‘bells and whistles’ of winning – appear to be interpreted as actual gains and skew the estimates of earned profits (Dixon et al., 2010; Dixon et al., 2015). An alternative possibility is that because the cues were not aligned with any particular behavioural goal, they may have distracted participants from the default risk-averse strategy of focusing on probabilities, thereby promoting risk. In either case, our findings lend support to the notion that sensory stimulation in gambling could act to de-emphasize the unfavourable odds of winning.

Pupillometry data pointed to additional and distinct effects. Firstly, we observed that riskier choices were associated with greater pupil dilation, independent of the presence of sensory cues. It is unlikely that the observed effects were driven by luminance, as the analyses only focused on visually identical trial periods. Although carry-over of feedback-related luminance effects could plausibly influence pupil response on the subsequent trial, this would only be relevant to trials preceded by wins. Our analysis found similar effects following 0-outcome trials, which had identical visual stimuli in the feedback period for both cued and uncued versions. Nor could the results be attributed to the effects of pupil foreshortening error from eccentric gaze: our analysis of fixations indicated that this would, if anything, produce the opposite effects on pupil size from the ones we saw. Our findings of strong relationships between pupil dilation and risk support, and extend, previous findings regarding pupil dynamics that accompany decision making. Several studies have examined pupil responses during decision making under uncertainty and observed that pupil dilation tracks the level of uncertainty (Satterthwaite et al., 2007; Lavín et al., 2013), is associated with surprise (Preuschoff et al., 2011), and temporally corresponds to the timing of decisions (Einhäuser et al., 2010). Our data highlight a strong association between pupil dilation and risky choice, which to our knowledge is a novel finding.

We also observed that sensory cues were associated with greater decision‐ and anticipation-related pupil dilation, an effect that was particularly evident following wins. This cue-driven amplification of pupil dilation points to arousal-promoting effects of sensory reward cues, which are distinct from their risk-promoting effects. As amplified pupil responses were evident not only during feedback, but during the decision and anticipation phases, cue-driven arousal appears to extend beyond circumscribed effects on feedback and to modulate the experience of the task more generally. Although our findings do not speak to the to the neural mechanisms of this effect, changes in pupil size are considered a proxy measure of noradrenergic signaling given the close correspondence between changes in pupil size and LC firing rates (Aston-Jones and Cohen, 2005; Murphy et al., 2014; Joshi et al., 2016). Therefore, cue-induced increases in pupil dilation could hypothetically reflect changes in LC-mediated noradrenergic signaling. Notably, LC firing rates have been theorized to modulate task engagement. The adaptive gain theory postulates that intermediate levels of tonic LC neuron firing and high phasic spiking facilitate exploitation of the task at hand, whereas high tonic LC activity and low levels of phasic spiking promote disengagement from the task and exploration of alternatives (Aston-Jones and Cohen, 2005). This could be relevant to disordered gambling, as playing modern electronic gambling machines that feature intense sensory stimulation has been associated with states of heightened engagement and immersion in problem gamblers, referred to as the “machine zone” or “dark flow” (Schüll, 2012; Dixon et al., 2014; Dixon et al., 2017). The hypothesis that sensory reward cues could promote immersion via noradrenergic modulation can be tested in future human and animal pharmacological challenge studies, as well as experiments using pupillometry. Indeed, changes in pupil size have been reported to co-vary with shifts between exploration-dominated and exploitation-dominated control states: exploration is associated with larger baseline pupil sizes and smaller phasic responses, whereas the opposite is seen during putatively exploitation-dominated states (Aston-Jones and Cohen, 2005; Gilzenrat et al., 2010; Jepma and Nieuwenhuis, 2011; Hayes and Petrov, 2016a).

We did not observe a cue-induced risk enhancement on the IGT, although participant performing the cued IGT were less likely to avoid risky decks immediately following large losses. The absence of a clear risk-promoting effect of cues on the IGT may be related to cues accompanying wins, whereas avoidance of risky decks on the IGT is largely driven by loss feedback. Though our findings appears to be at odds with the findings on the cued Rodent Gambling Task (Barrus and Winstanley, 2016), there are important differences between the human and rodent tasks. The human IGT assesses not only risk, but also learning from rewards and punishments. Rats undergo extensive training to ensure development of stable, asymptotic choice preferences, and also learn reward probabilities experientially *before* the actual testing through equivalent numbers of forced choice trials per “deck”. In this regard, the rodent task could be considered more similar to the VGT, where the probabilities are known rather than gradually learned.

Our study had several limitations. First, pupillometry analysis was not performed on the IGT due to the challenge of defining a ‘baseline’ period given the temporal structure of the task. Testing in two eye-tracking laboratories using slightly different equipment entailed our VGT pupillometry analysis be restricted to a subset of participants, but we corroborated the behavioural effects within that subsample. Although we limited our analysis to visually identical phases of the two VGT versions, carry-over luminance effects from feedback cannot be ruled out, although as mentioned earlier, we consider this possibility unlikely. A significant limitation is the lack of correction for pupil foreshortening error resulting from gaze eccentricity, which distorts pupil area measures, as the pupil assumes a more elliptical shape, as registered by the camera, at greater eccentricities. To avoid this issue, many pupillometry experiments require central fixation, which is not optimal from the ecological standpoint. As mentioned earlier, our analysis of gaze positions during the relevant time intervals revealed only non-significant differences in fixation eccentricity between cued and uncued VGT versions, which would be expected, if anything, to drive pupil size in the opposite direction of our findings.

In conclusion, we found that sensory reward-paired cues can promote riskier choice in healthy human volunteers. To our knowledge this is the first direct demonstration of risk-promoting effects of such cues in human subjects. We also observed effects of these stimuli on pupil dynamics that were independent of their risk-promoting effects. Rather, they appeared pertain to task experience, be it global task-related arousal or more specific changes in LC-mediated control states, which could promote maladaptive task engagement. Currently, there is no regulation around the integration of sensory cues into commercial gambling products. Both these observations are consistent with the view that the presence of such cues in commercial gambling products may facilitate problematic gambling behaviour.

## Acknowledgments

We thank D. Griffin, R. Gonzalez and E. Limbrick-Oldfield for consultation on data analysis.

Funding: This work was supported by National Center for Responsible Gaming (NCRG) seed funding awarded to MVC and research allowance funds from post-doctoral fellowships from CIHR and the Michael Smith Foundation for Health Research by which MVC was supported. This work was also partially supported by an operating grant awarded to CAW, LC and AJS from the Canadian Institutes for Health Research (CIHR; PJT-056012). MVC has received a speaker honorarium from the Responsible Gaming Association of New Mexico. In the past three years, CAW has consulted for Shire and Hogan Lovells LLP and received due compensation. LC is the Director of the Centre for Gambling Research at UBC, which is supported by funding from the Province of British Columbia and the British Columbia Lottery Corporation (BCLC), a Canadian Crown Corporation. The Province of British Columbia government and BCLC had no involvement in the ideas expressed herein, and impose no constraints on publishing. LC receives funding from the Natural Sciences and Engineering Research Council (Canada). LC has received a speaker honorarium from Svenska Spel (Sweden) and an award from the NCRG (US). He has not received any further direct or indirect payments from the gambling industry or groups substantially funded by gambling. He has provided paid consultancy to, and received royalties from, Cambridge Cognition Ltd. relating to neurocognitive testing. The authors confirm they have no other conflicts of interest or financial disclosures to make.

